# LiveLattice: Real-time visualization of tilted light-sheet microscopy data using a memory-efficient transformation algorithm

**DOI:** 10.1101/2024.05.28.596280

**Authors:** Zichen Wang, Hiroyuki Hakozaki, Gillian McMahon, Marta Medina-Carbonero, Johannes Schöneberg

## Abstract

Light-sheet fluorescence microscopy (LSFM), a prominent fluorescence microscopy technique, offers enhanced temporal resolution for imaging biological samples in four dimensions (4D; x, y, z, time). Some of the most recent implementations, including inverted selective plane illumination microscopy (iSPIM) and lattice light-sheet microscopy (LLSM), rely on a tilting of the sample plane with respect to the light sheet of 30-45 degrees to ease sample preparation. Data from such tilted-sample-plane LSFMs require subsequent deskewing and rotation for proper visualization and analysis. Such transformations currently demand substantial memory allocation. This poses computational challenges, especially with large datasets. The consequence is long processing times compared to data acquisition times, which currently limits the ability for live-viewing the data as it is being captured by the microscope. To enable the fast preprocessing of large light-sheet microscopy datasets without significant hardware demand, we have developed WH-Transform, a novel GPU-accelerated memory-efficient algorithm that integrates deskewing and rotation into a single transformation, significantly reducing memory requirements and reducing the preprocessing run time by at least 10-fold for large image stacks. Benchmarked against conventional methods and existing software, our approach demonstrates linear scalability. Processing large 3D stacks of up to 15 GB is now possible within one minute using a single GPU with 24 GB of memory. Applied to 4D LLSM datasets of human hepatocytes, human lung organoid tissue, and human brain organoid tissue, our method outperforms alternatives, providing rapid, accurate preprocessing within seconds. Importantly, such processing speeds now allow visualization of the raw microscope data stream in real time, significantly improving the usability of LLSM in biology. In summary, this advancement holds transformative potential for light-sheet microscopy, enabling real-time, on-the-fly data processing, visualization, and analysis on standard workstations, thereby revolutionizing biological imaging applications for LLSM, SPIM and similar light microscopes.

## INTRODUCTION

### A tilted objective angle in light-sheet microscopy leads to unique preprocessing challenges

Light-sheet fluorescence microscopy (LSFM) is a principal type of fluorescence microscopy extensively employed in cell and developmental biology^1–10^. Compared to point-scanning confocal microscopy, LSFM offers significantly enhanced temporal resolution and duration for three-dimensional time-lapse imaging (4D; x, y, z, time) by simultaneously illuminating and capturing images across an entire plane.

While there are multiple different configurations for how the light sheet and the perpendicular detection objective are realized^11^, one major distinction arises from how the two-objective configuration is oriented with respect to the sample. While a 0°angle relative to the sample holder is common, tilted angles are often adopted resulting in tilted-sample-plane LSFMs. Examples are 45° angles in inverted selective plane illumination microscopy (SPIM)^12^ and open-top SPIM^13^, or 30° angles in lattice light-sheet microscopy (LLSM)^14,15^ (**Figure 1A**).

**Figure 1.**
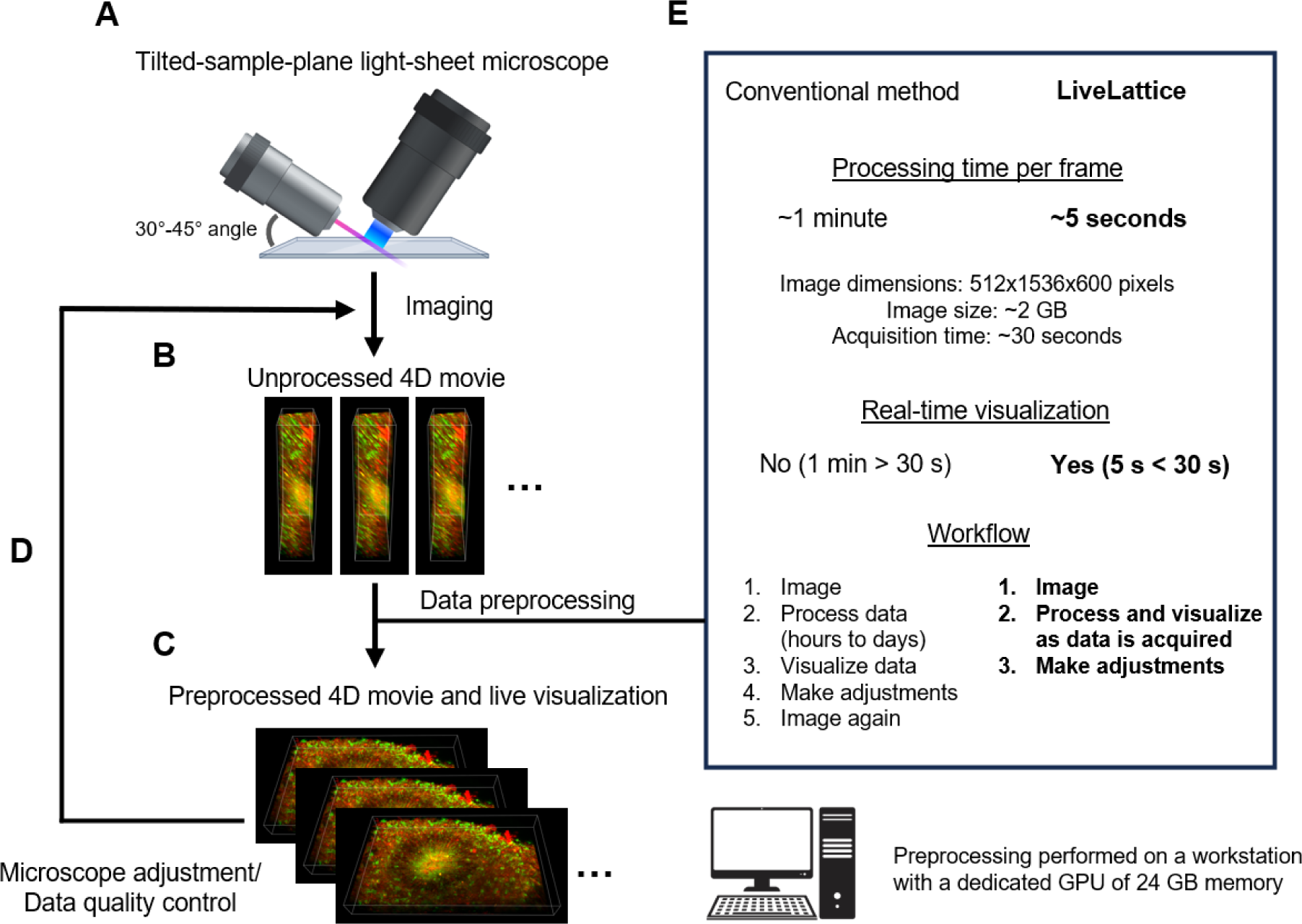
Ten-fold acceleration of tilted-sample-plane light-sheet microscopy preprocessing allows real-time data visualization. (A) The combined excitation/detection objectives are tilted with respect to the sample plane. E.g. by ∼30° in lattice light-sheet microscopy. (B) Raw 3D image stacks from such microscopes are skewed and require preprocessing before visualization. (C) After data preprocessing, a 4D movie for the region of interest is revealed. (D) Microscope adjustment and data quality control (QC) steps can now be performed to inspect whether the region of interest is desirable and whether the imaging parameters need to be adjusted (E) Current preprocessing algorithms are too computationally expensive to allow real-time visualization of large 4D dataset, leading to a workflow that decouples imaging from data QC. Here, we proposed a novel memory-efficient algorithm, WH-Transform, that leads to a 10-fold acceleration in preprocessing speed and thereby allows real-time 4D data visualization and QC.

To achieve volumetric imaging in these configurations, the sample is moved laterally to acquire a 3D stack of skewed images. This skewed 3D stack cannot be readily viewed or processed (**Figure 1B**). Instead, the stack must undergo a coordinate transformation preprocessing step in order to be visualized and analyzed (**Figure 1C**).

Recent implementations of tilted-sample-plane LSFM, such as LLSM, produce large skewed image stacks, often exceeding multiple gigabytes per volumetric stack. Preprocessing these stacks typically involves deskewing, axis scaling, and rotation. Current preprocessing methods, for example cudaDecon^16^, require substantial memory and often employ data cropping to accelerate preprocessing speed.

Large multi-dimensional image transformation calculations are most efficiently done in a graphic processing unit (GPU), as it has 2-3 orders of magnitude more cores than a central processing unit (CPU)^17^. However, a GPU is limited by the on-board dedicated memory size. When the memory requirements of a processing task exceed the GPU limit, memory shuffling between GPU and CPU can create excessive overhead that drastically increases the computation time^18^. Alternatively, high performance computing (HPC) clusters^19^ can offer up to a few hundred GBs of memory, but are less accessible and flexible for routine data preprocessing, or real-time visualization. Large volumetric data can also be processed using CPU which usually comes with up to several hundred GBs of memory, but is considerably slower as mentioned. Thus, in order to process large data volumes, the data is subdivided, processed, and combined again using image stitching^20^. However, the additional stitching step further increases the computation time and may introduce artifacts due to misalignment.

As a result, image stacks that take under a minute to acquire using tilted-sample-plane LSFM may take often hours to be preprocessed using accessible hardware. One of the most impactful problems from this lag is the inability for live visualization of the sample as it is being imaged. Live sample visualization and data inspection during acquisition is a crucial feedback mechanism for microscopy (**Figure 1D**). Live inspections might include screening for suitable regions of interest, re-centering the stage, or focusing adjustments, none of which are feasible with current tilted-sample-plane LSFM techniques such as LLSM. As a consequence, microscopy data sometimes has to be discarded after acquisition because a flaw is revealed in the visual post hoc inspection. Ideally, tilted-sample-plane LSFM data could be preprocessed in a way that allows live visualization of the data as it is being acquired.

Here, we introduce LiveLattice, a fast data preprocessing pipeline for tilted-sample-plane LSFM. LiveLattice utilizes a novel Wang-Hakozaki Transformation (WH-Transform) algorithm to perform memory-efficient deskewing and rotation of the data. Using WH-Transform, LiveLattice drastically cuts memory demands and processing times for data from such microscopes. For a 2 GB image stack, our method is nearly 10-fold faster, enabling live visualization of 4D data as it streams from the microscope (**Figure 1E**).

## RESULTS

### A combined deskewing and rotation algorithm drastically reduces memory requirements for skewed 3D image stack preprocessing

Skewed 3D stacks that are the result of tilted-sample-plane LSFM require a three-step process before visualization and further processing (**Figure 2A**).

**Figure 2.**
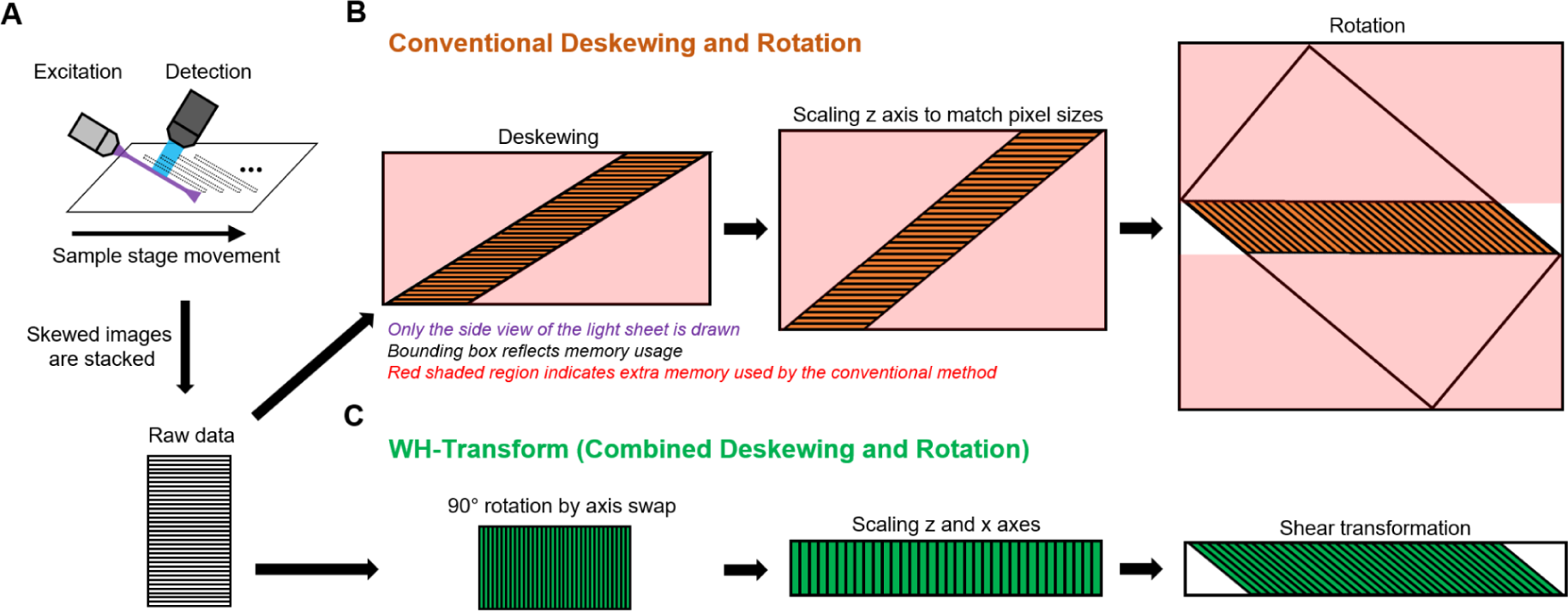
Novel combined transformation drastically reduces memory usage compared to the conventional method. (A) Top: Tilted-sample-plane LSFMs illuminate the sample at a tilted angle relative to the sample stage motion (top). Bottom: The resulting raw data is a skewed 3D image stack. Only the side view of the light sheet is shown. (B) Conventionally, the 3D stack is deskewed, isotropically scaled, and rotated to transform the image into the sample coordinates. However, in order to perform the computation, large padded regions are allocated in the memory (black box indicates full memory size and red shaded region denotes unnecessary memory usage). (C) To reduce memory allocation during preprocessing, we combined deskewing, scaling, and rotation into a memory-efficient three-step operation (WH-Transform). First, 90° rotation along the y-axis is done by swapping the other two axes. Second, we scaled the axes other than y to match the dimensions of the final processed volume. Third, we performed a shear transformation to fully transform the volume into the sample coordinates.

First, the raw image stack is deskewed to shift each plane and recapitulate the sample geometry. Second, the stack is isotropically scaled to match the lateral and axial pixel sizes. Third, the deskewed and scaled image stack is rotated so that the vertical axes in the data and the sample are aligned (**Figure 2B**).

Current implementations of this three-step process, such as cudaDecon, use zero-value pixels as padding in every step (red shaded region in **Figure 2B**) to facilitate deskewing, scaling, and rotation. As the imaged sample volume increases, the memory size required for deskewing and rotation operations increases polynomially. For example, for a 512×1536×800 stack, around 90% of the volume used for rotation is used as padding. These empty pixels lead to a very large memory overhead that significantly slows down computation time, even on the most efficient hardware using the most accelerated implementations.

To enable the fast preprocessing of large light-sheet microscopy datasets without significant hardware demand, we have developed WH-Transform, a novel light-sheet preprocessing algorithm. Instead of sequentially performing deskewing and rotation which requires considerably more memory usage than the processed data (bounding boxes around the data in **Figure 2B**), we combine both operations and thereby avoid the need for any extra memory beyond the size of the processed volume (bounding boxes around the data in **Figure 2C**).

To avoid the use of a large memory allocation, we designed a different series of image transformation operations to reach the same deskewed and rotated volumes. First, we rotated the image by 90° along the sample y-axis by swapping the 3D array axes. This step essentially takes no run time, yet proves useful for processing volumes with a large number of planes. Second, we performed a scaling transformation in two axes to account for 1) differences in lateral and axial pixel sizes, and 2) actual sample thickness after deskewing. Third, we incorporated the effects of both deskewing and rotation into a single shear transformation. As is evident in **Figure 2C**, no additional padding pixels are required, and the only empty pixels are present around the corners where the light sheet meets the sample at a tilted angle.

### WH-Transform leads to linear preprocessing and a 10-fold acceleration over conventional methods

We benchmarked our WH-Transform algorithm against two light-sheet data deskewing and rotation methods for a range of raw data sizes using one NVIDIA RTX 3080 Ti with 24 GB memory.

Three example datasets were used with endogenously tagged Tom20-EGFP (mitochondria, green) and CAAX-RFP (membrane, red) acquired using a custom-built lattice light-sheet microscope (see Methods): A 0.7 GB 512×1536×200 px raw 3D stack of adherent hepatocytes **(Figure 3A, circle)**, a 2.1 GB 512×1536×600px raw 3D stack of a human cortical brain organoid **(Figure 3B, triangle)**, and a 3.4 GB 512×1024×1500 px raw 3D stack of a human branching lung organoid **(Figure 3C, square)**.

**Figure 3.**
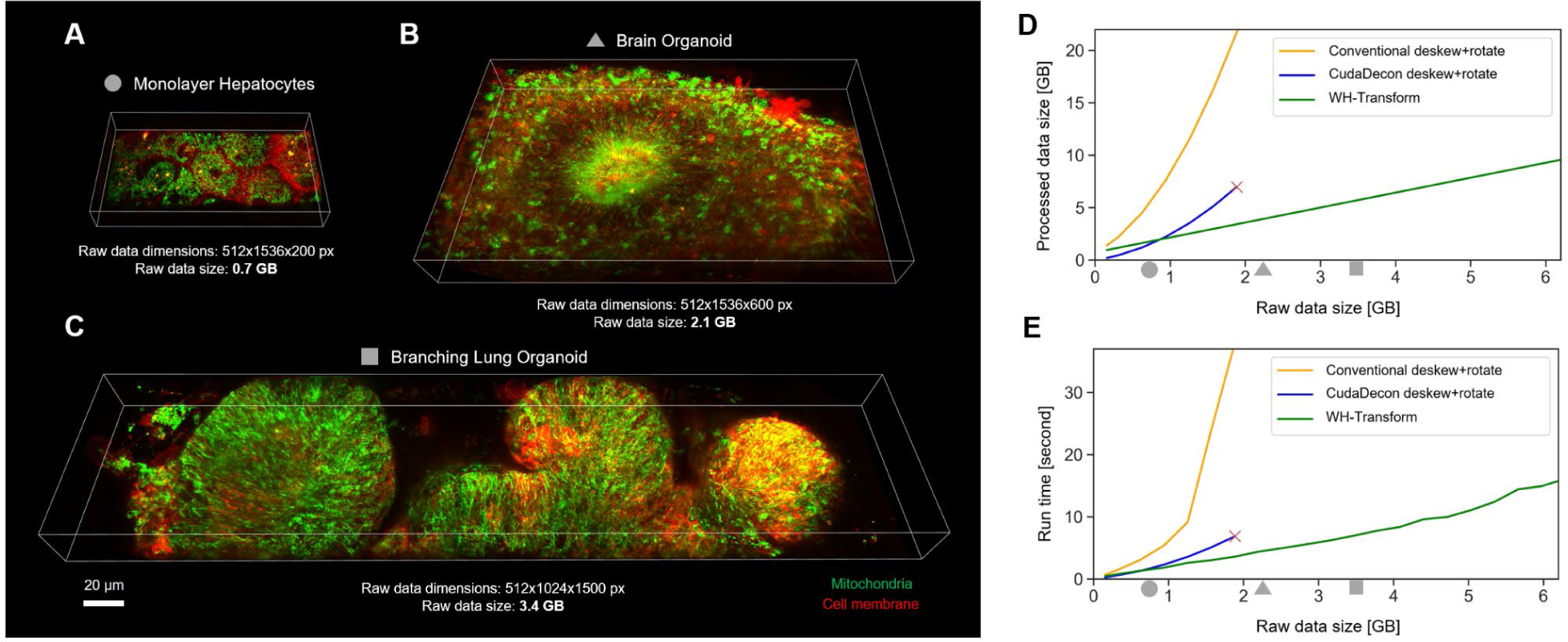
Benchmark of light-sheet data preprocessing methods shows that WH-Transform achieves linear scaling and drastically faster speed for larger dataset. (A-C) Example processed lattice light-sheet data for three increasingly larger raw data sizes. A: monolayer hepatocytes with raw data size 0.7 GB, B: human cortical brain organoid rosette with raw data size 2.1 GB, C: human branching lung organoid with raw data size 3.4 GB. Mitochondria are shown in green and plasma membranes are shown in red. (D-E) Processed data size (D) and run time (E) as a function of input raw data. Orange: Conventional deskewing and rotation implemented using GPU-based affine transformation in Python. Blue: cudaDecon software commonly used for lattice light-sheet data preprocessing. Red cross indicates the software was not able to process larger datasets using the same hardware. Green: WH-Transform that combines deskewing and rotation to avoid unnecessary memory allocation. Gray symbols indicate where the example datasets in A-C sit on the raw data size axis.

We first implemented the conventional deskewing and rotation methodology using GPU-based affine transformation (**Figure 3D, E, orange curves**). Both the processed file size (**Figure 3D**) as well as the run time (**Figure 3E**) increase geometrically as the size of the input raw data increases linearly. Larger processed data sizes considerably slow down the overall computation mainly due to inefficient GPU-CPU data shuffling. Next, we tested the popular GPU-accelerated preprocessing package cudaDecon^16^ (**Fig 3D, E, blue curves**). By cropping the rotated volume to only the region where there appears to be sample data, cudaDecon is able to significantly reduce the processed file size and run time. However, the sample cropping may lead to suboptimal results where important regions of the data are removed or extra padding is added to larger datasets. Additionally, in our hands on our system, cudaDecon was not able to process raw 3D stacks that are larger than 2 GB due to memory overflow (24 GB GPU memory) (**Fig 3D, E, red crosses**). Note that both datasets in **Figure 3B, C** are too large to be processed using cudaDecon and would have to be split into multiple sub regions in order to be processed. Finally, we benchmarked WH-Transform, our combined deskewing and rotation algorithm (**Fig 3D, E, green curves**). WH-Transform 1) processes the original data without image cropping, 2) leads to linear scaling in both file size and runtime as raw data sizes increase, 3) is able to process much larger files compared to the cudaDecon limit using the same set of hardware, and 4) leads to a run time that is at least 10 times faster than the conventional method for large volumes.

### WH-Transform allows for real-time visualization of large light-sheet volumes

Next, we compared how existing methods for preprocessing compare to WH-Transform in a live/real-time visualization scenario. We integrated the WH-Transform into our custom LLSM data deconvolution and preprocessing pipeline and tested it on a human cortical brain organoid sample with endogenously tagged Tomm20-EGFP (mitochondria, green) and CAAX-RFP (membrane, red). The sample was imaged in 3D using a 512×1536 camera size for 600 scans at an exposure time for each scan of 50ms, summing up to 30s acquisition time per 3D volume. The raw image data is shown in **Figure 4A**. The preprocessed volume is shown in **Figure 4B**.

**Figure 4.**
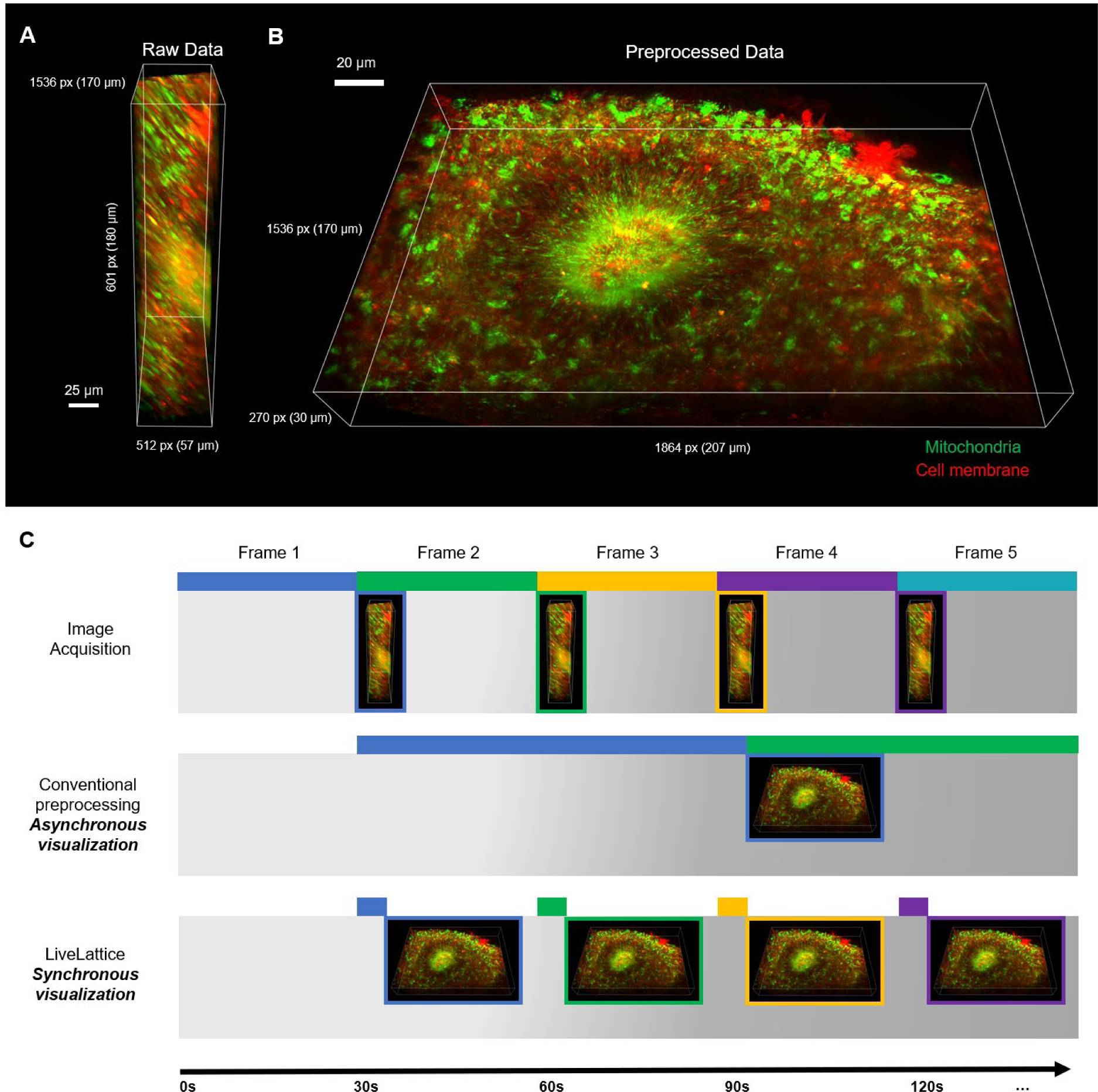
Demonstration of WH-Transform-based real-time visualization of a 4D brain organoid LLSM dataset. A) Representative raw data with dimensions 512×1536×800 pixels. Mitochondria are shown in green and plasma membrane is shown in red. B) The raw data is deconvolved and transformed using WH-Transform. The processed data has dimensions 2492×1536×270 pixels or 276×170×30 microns. C) Illustration and comparison of conventional method and LiveLattice for live visualization of 4D dataset. Top: raw image stack is acquired every 30 seconds on the microscope. Middle: preprocessing based on conventional algorithm required more than 70 seconds to process one stack, resulting in significant delays between data acquisition and visualization. Bottom: preprocessing based on WH-Transform required around 5 seconds, thus each timepoint can be processed before the next frame is acquired, enabling synchronous visualization.

In our hands, cudaDecon was not able to process the data due to insufficient memory availability using our 24 GB GPU. The affine transformation implementation took ∼70 seconds to process a single volumetric frame, moving the availability of frame 1 for visual inspection to after the acquisition of frame 4 (**Figure 4C, middle row**). In contrast, the WH-transform’s combined deskewing and rotation transformation finished the preprocessing after about 5 seconds, making the first timepoint available for visualization before the acquisition of the second timepoint was completed (**Figure 4E**).

### LiveLattice: Implementation of WH-Transform in a lattice light-sheet preprocessing pipeline makes real-time visualization available to the LLSM community

As a last step, we incorporated WH-Transform-based deskewing and rotation into our routine lattice light-sheet data processing pipeline to create LiveLattice. First, deconvolution is performed directly on the skewed dataset using a point spread function (PSF) acquired in sample scan mode as described recently^21^. The advantage of this approach includes 1) more accurate deconvolution because the PSF is acquired in the same way as the data by scanning the light sheet along the sample stage, and 2) faster deconvolution because the size of the raw data is smaller than the deskewed and rotated data. For this deconvolution, we first used the python wrapper of cudaDecon, pycudaDecon, while omitting its deskew and rotate functionality. Second, we then applied WH-Transform to efficiently deskew and rotate the deconvolved image stack. To preprocess large 3D image stacks most effectively, both deconvolution and deskewing/rotation steps utilize GPU-based parallel-computing through CUDA^22,23^ and Dask^24^. As data streams from the microscope, our pipeline detects newly acquired stacks and automatically preprocesses them.

Once the data has been preprocessed, it can be loaded into visualization software **(Figure 5B)**. We use Napari^25^ to live view our lattice light-sheet data as it streams from the microscope. To make this possible for files that exceed local memory, we used delayed data loading based on Dask^24^. Besides data visualization, further downstream data processing/analysis can be performed on the preprocessed 4D dataset **(Figure 5C)**. In the case of 4D particle tracking, pyLattice ParticleTracking^26^ can be applied for particle segmentation and tracking. In the case of 4D analyses of mitochondrial motility and dynamics, pyLattice MitoTNT^27^ can be used to reliably track and extract functional information from mitochondrial networks.

**Figure 5.**
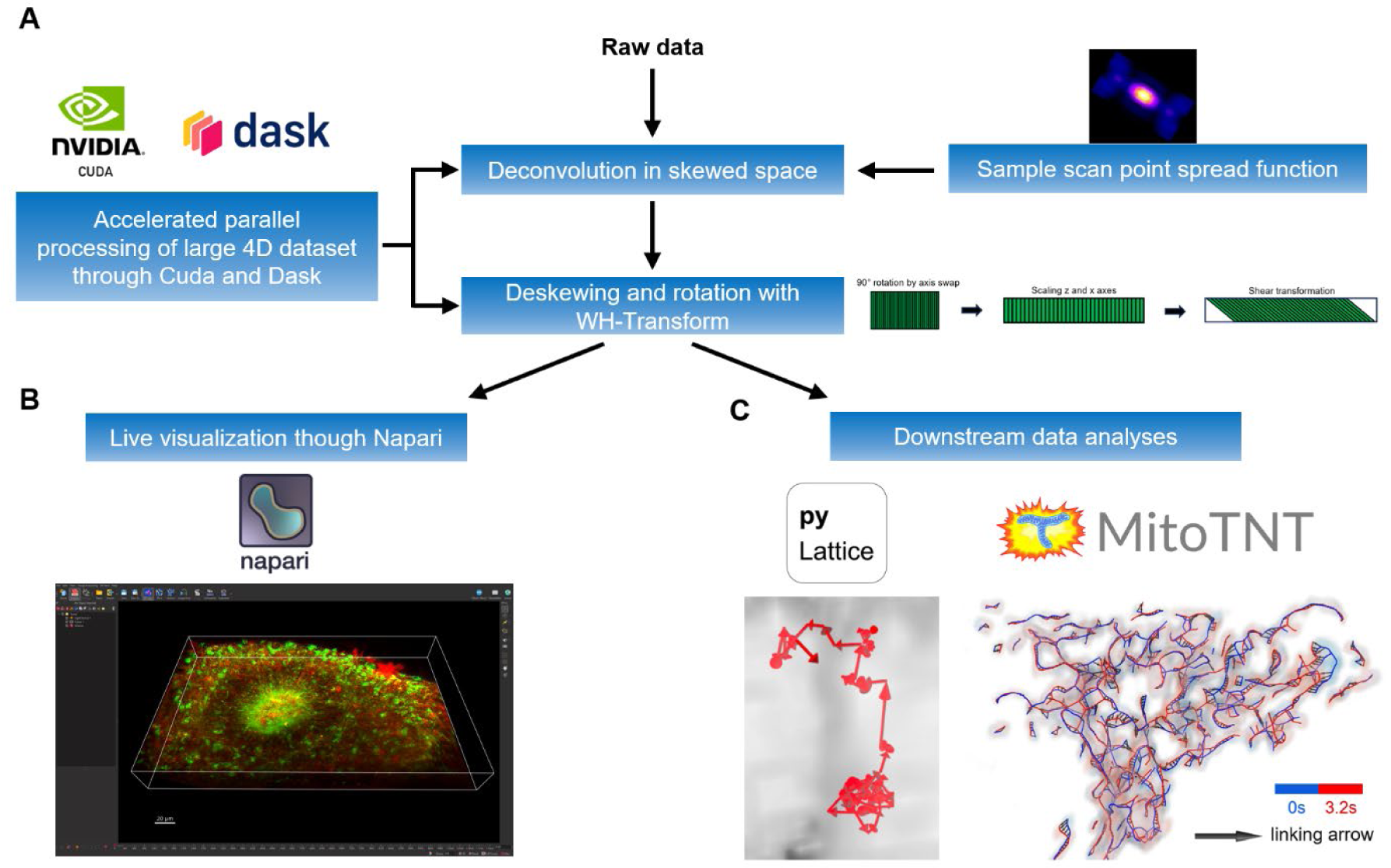
LiveLattice, a GPU-based parallel-computing pipeline for large 4D lattice light-sheet data preprocessing, live visualization, and processing. Raw data is processed in the following steps: A) First, deconvolution is performed by cudaDecon on the skewed dataset using a sample scan point spread function (PSF). Second, deskewing and rotation is performed using WT-Transform. To efficiently preprocess large 3D image stacks, both deconvolution and deskewing/rotation utilize GPU-based parallel-computing through CUDA and Dask. B) Preprocessed data can then be dynamically loaded into a visualization software such as Napari for real-time visualization during imaging sessions, and C) be used for further downstream data analyses such as for particle tracking (pyLattice ParticleTracking) or organelle tracking (pyLattice MitoTNT).

To enable real-time viewing of lattice light-sheet data during acquisition, we have integrated LiveLattice into pyLattice, a Python library for advanced lattice light-sheet image analysis. It can be accessed at https://github.com/pylattice/liveLattice.

## DISCUSSION

Light-sheet fluorescence microscopy has revolutionized biological imaging by offering four-dimensional (4D; x, y, z, time) data with high spatial and temporal resolution. Many recent light-sheet microscope implementations such as lattice light-sheet microscopy illuminate the sample at a tilted angle, which necessitates image preprocessing through deskewing and rotation for proper visualization and further processing. However, the current algorithms for deskewing and rotation of large light-sheet data require significant memory allocations, which makes the GPU-based computations time-consuming. To allow fast preprocessing of 4D light-sheet datasets, we have designed a memory-efficient transformation algorithm that performs remarkably better (∼10-fold better) on large light-sheet datasets compared to the existing software. Instead of performing deskewing and rotation separately, we combined both operations into a series of affine transformations that we call WH-Transform. By avoiding superfluous memory usage, the combined transformation drastically reduces preprocessing time, file storage space, as well as the need for expensive hardware or HPC resources. The capability to rapidly preprocess large 3D volumes also reduces or even eliminates the need for image stitching that was previously required for imaging large fields of view. Importantly, combined with fast deconvolution and visualization (e.g. through Napari), our GPU-accelerated memory-efficient preprocessing method has enabled the real-time visualization of large 4D light-sheet dataset on a regular workstation. This on-the-fly preprocessing and visualization pipeline that we call LiveLattice can revolutionize the field of light-sheet microscopy, and better the way for researchers to use these advanced microscopy methods to study important questions in biology and human health.

## METHODS

### Sample Preparations

#### Stem Cells

Both brain and lung organoids were differentiated from a human induced pluripotent stem cell (hiPSC) line tagged with CAAX-RFP for the plasma membrane and Tom20-EGFP for mitochondria. An Tom20-EGFP-tagged mitochondriacell line was obtained from the Allen Institute for Cell Science through the UCB Cell Culture Facility (AICS-0011 cl.27). Into this cell line, the CAAX domain of K-ras tagged with mTagRFP-T was introduced into a safe harbor locus via CRISPR using a donor plasmid developed at the Allen Institute for Cell Science and sourced from Addgene (Plasmid #107580a). We call the resulting cell line CT20. CT20 hiPSCs were cultured using standard hiPSC tissue culture techiques and underwent periodic testing for mycoplasma and pluripotency.

### Human Hepatocytes

Monolayer hepatocytes were differentiated from CT20 hiPSCs using STEMCELL Technologies’ STEMdiff™ Hepatocyte Kit. For days 1-4, hiPSCs were directed towards definitive endoderm. For days 5-10, definitive endoderm cells were differentiated into hepatic progenitor cells. After another 10 days, hepatic progenitor cells were matured into hepatocyte-like cells.

### Human Lung organoids

Branching lung organoids (BLOs) were prepared according to STEMCELL Technologies’ STEMdiff Branching Lung Organoid kit (#100-0195). Briefly: Prior to differentiation, CT20 hiPSCs were clump passaged and grown to 30-60% confluency in a 24-well plate. The definitive endoderm was formed as a dense monolayer of cells (day 0-3). Anterior foregut endoderm buds floated into the media from the monolayer (day 3-6). These were removed and further differentiated into lung bud organoids free-floating in media (day 6-14). Lung bud organoids were differentiated into BLOs in a Matrigel (Corning, 356231) sandwich within a Transwell insert (day 14-42+).To prepare for imaging with LLSM, the Matrigel sandwich was dissociated and the BLO was washed. It was resuspended in Matrigel and pipetted onto a coverglass in a dome. BLOs were imaged in phenol red-free DMEM/F12 (Gibco, 11320033) supplemented with 2% FBS (Genesee Scientific 25-514H).

### Human Brain organoids

Brain organoids were differentiated using a STEMCELL Technologies kit (Catalog #08570), following the manufacturer’s instructions. Briefly, embryoid bodies (EBs) were formed by placing 9,000 hiPSCs into a U-bottom 96-well plate (day 0) to promote germ layer differentiation. At day 5, the EBs were transferred into a 24-well plate to induce neural ectoderm differentiation. By day 7, the organoids were embedded with Matrigel (Corning, 356231) to provide external support and promote the expansion of neuroepithelial buds. By day 10, the organoids were transferred into a 6-well plate and placed on a shaker (75 rpm) to promote brain tissue growth and expansion. At day 30, the organoids were cultured in Neurobasal media (Gibco, 21-103-049) containing 2% B27 (Gibco, 17504044), 0.5% (wt/vol) anhydrous glucose, and 1% GlutaMAX (Thermo Fisher Scientific, 35050038). Brain organoids were imaged with the same culture media but without phenol red ((Neurobasal media PRF; Gibco, 17504044), 2% B27, 0.5% (wt/vol) anhydrous glucose, and 1% GlutaMAX).

### Lattice Light Sheet microscopy

A custom built lattice light sheet microscope designed by Eric Betzig’s Lab at HHMI Janelia/UC Berkeley was used to image samples. Key modifications of this work are the use of 0.6 NA excitation objective lens (Thorlabs, TL20X-MPL), a 1.0 NA detection objective lens (Zeiss W Plan-Apochromat 20×/1.0, model #421452-9800), and a Hamamatsu Photonics Orca Fusion BT sCMOS Camera for image acquisition. 488nm and 560nm lasers were used to excite GFP and RFP. A Multiple Bessel Beam Light Sheet Pattern with NA Max 0.4, NA Min 0.35 was used for the organoid samples and a Hexagonal Lattice Light Sheet Pattern with NA Max 0.5 and NA Min 0.42 was used for the monolayer cell samples. A 50msec exposure time was used per 2D scan and samples were scanned every 300nm continuously with a piezo-actuated stage. The three biological samples imaged (hepatocytes, brain organoid, and lung organoid) were scanned for 200, 600, 1500 slices, or 60, 180, 450 microns respectively. The volumetric frame rates were 10, 30, and 75 seconds per frame, respectively.

### Benchmark Hardware

We used a custom-built workstation with an AMD® Ryzen 9 5950×16-core processor, an NVIDIA RTX 3080 Ti 24 GB GPU, 128 GB memory, and Ubuntu 20.

## ACKNOWLEDGEMENTS

This study was supported by funds from the Hartwell Foundation through an Individual Biomedical Research Award to J.S., through an NIH Director’s New Innovator Award to J.S., through an W.M. Keck Award to J.S., and through an NIH NIBIB Award to G.M. under Award number T32EB009380. The MitoTNT logo was created by Andre Modolo in the Schöneberg lab.

## AUTHOR CONTRIBUTIONS

### Authors and Affiliations

All authors were part of the Department of Pharmacology, University of California, San Diego, San Diego, CA, 92093 and the Department of Chemistry and Biochemistry, University of California, San Diego, San Diego, CA, 92093.

### Contributions

Hiroyuki Hakozaki (HH) conceived the algorithm. Zichen Wang (ZW) implemented and benchmarked the algorithm, performed the human hepatocyte culture and sample preparation. Gillian McMahon (GM) performed the human lung organoid culture and sample preparation. Marta Medina Carbonero (MMC) performed the human cortical brain organoid culture and sample preparation. Johannes Schöneberg (JS), ZW, HH, GM, and MMC wrote the manuscript. JS was responsible for conceptualization, funding, and administration.

### Corresponding author

Correspondence to Johannes Schöneberg.

## DECLARATION OF INTERESTS

UCSD has filed for patent protection on the technology described herein.

## Notes

https://github.com/pylattice/LiveLattice

### Competing Interest Statement

The authors have declared no competing interest.

